# A self-similar helix as a metaphor for continuous time with nested periodicity

**DOI:** 10.1101/2025.03.06.640619

**Authors:** Richard Bischof

## Abstract

Humans perceive time as both linear and circular, which is reflected in the dichotomous way we typically organize and visualize temporal information in science: as either time series or periodic plots. These seemingly incompatible perspectives can be reconciled with a helical representation of time. Helices have been underused as a visualization of temporal information, in part because two-dimensional visualization of three-dimensional structures leaves portions of the data hidden and in part due to inherent disadvantages of radial visualizations over linear ones. Nonetheless, the helical metaphor of time brings along an important advantage - it offers an intuitive way to depict continuous time with periodicity at multiple scales. Here, I introduce a representation of time via a model of self-similar, or hierarchical, helices. This model can serve as a cognitive map or mental image of time, as well as an immersive data visualization approach for exploring patterns in timed data with nested periodicity. I demonstrate the model with a few examples (astronomical light phases, air temperature profiles, and wildlife activity patterns) and provide R functions to generate interactive displays of temporal data projected onto hierarchical helix structures.

## Main text

### Reconciling linear and circular representations of time

Time, the natural phenomenon and fundamental quantity, may or may not exist as a flow^1^. Yet, our conceptual construct of time as such helps us organize our lives and catalogue the dynamics of the world around us. This construct varies substantially between and even within cultures^2^, but there is one recurring theme: people perceive time’s progression as both linear and circular^3^. This dichotomy is also manifested in the way we visualize timed information in science, where time series and periodic plots are the most common methods for displaying temporal data.

Although the linear and circular concepts of time seem incompatible^4^, they can be reconciled by a helical representation, with loops of the helix representing periodic elements and the direction in which the helix stretches representing time’s forward progression (Fig. 1). This conceptual model has occasionally been put forth as a way of visualizing temporal data^5–7^, including for spatio-temporal applications^8^. As a metaphor for time, the helix also found its way into psychology^9^,^10^. A two-dimensional version of the three-dimensional time helix is the spiral graph^6^,^11^,^12^. The helical or spiral perspectives on time also feature in art and were among the self-reported perceptions during an extensive online public survey conducted in Norway in 2017^4^.

**Figure 1:**
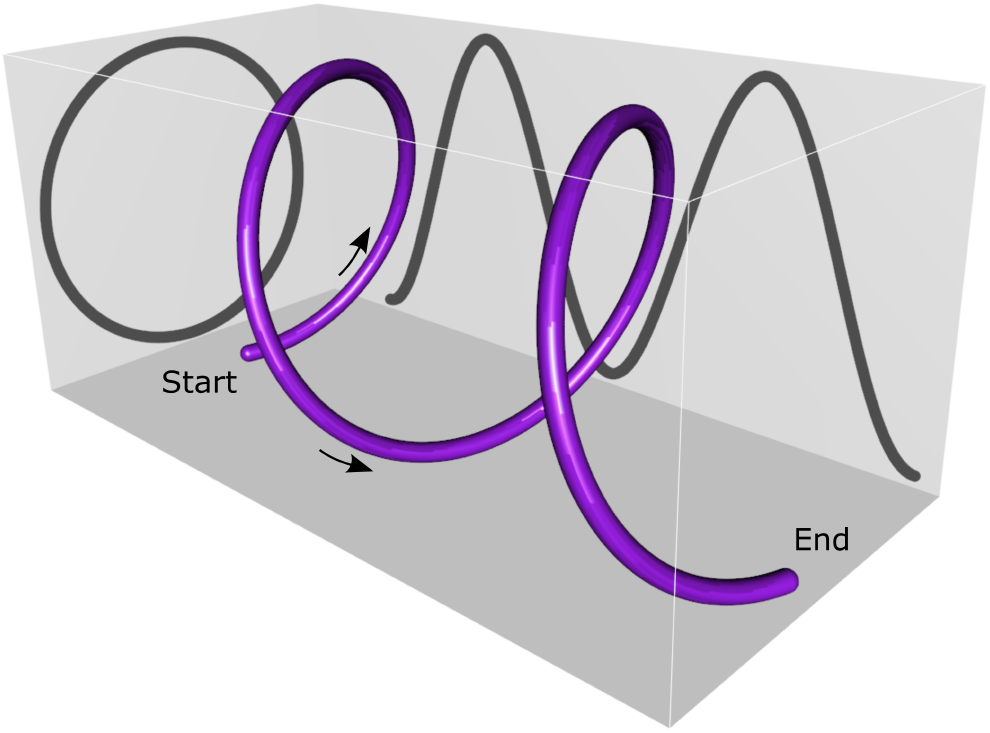
A helical representation of time reconciles the linear and circular perspectives, represented by the shadows projected onto the vertical planes of the background. The arrows indicate the direction of the passage of time along the helical structure.

As an example, we can let each loop (referred to as “round” by^13^) of a helix represent a 24-hour day and the succession of loops the progression of time over multiple days. This allows visualization of daily phenomena, such as alternating periods of daylight and dark, as patterns aligned along the longitudinal axis of the helix. Simultaneously, it can reveal changes in periodicity in observations over a longer duration, like the lengthening and shortening of the period of daylight across seasons at northern latitudes (Figure 2).

**Figure 2:**
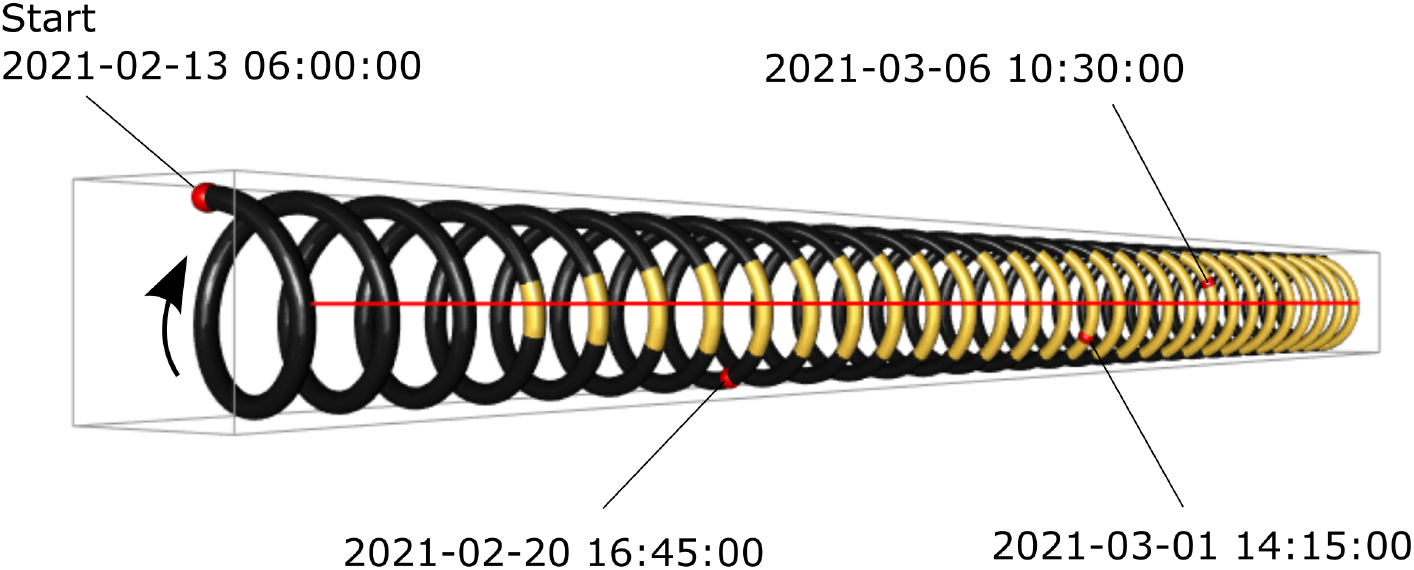
Time helix with 24-hour loops over a period of 30 days. Colors indicate daylight (yellow) and dark (black) based on civil sunrise and sunset hours at Longyearbyen on the island of Svalbard (15.47^*?*^ E, 78.25^*?*^ N) around the end of polar night in February 2021. The red line marks noon; example dates and times are shown with red dots. The arrow indicates the direction along which time passes.

### Hierarchical helix and periodicity at multiple scales

Although intuitively appealing, the helical representation of time remains uncommon. This is in part because, with a three-dimensional construct projected into a two-dimensional image, users may not see full cycles^6^ and in part because information is more readily decoded and compared when represented in linear instead of radial form^14^. Furthermore, the primary information shown with a single helix in Fig. 2 is easily conveyed by mapping it onto two dimensions (for an example, see^15^), albeit with the loss of visualized periodicity. Nevertheless, the helical model of time holds an unexploited potential, lacking or limited in other representations: it can be readily extended to allow a cognitive map of periodicity at multiple temporal scales simultaneously, without breaking the time continuum.

Many phenomena, including the lives of humans and other organisms, take place over, and are influenced by, nested periodic events. For example, the dynamics of life on earth are to a large extent governed by the rotation of the planet around its axis (1 earth day = 24 hours) and the accompanying cycles in daylight and temperature; lunar orbits (1 lunar month ≈ 29.5 earth days) and the resulting tidal and nocturnal illumination patterns; and earth’s revolution around the sun (1 earth year ≈ 365.24 earth days) and associated seasonal changes. Astronomical events from a geocentric perspective are only one example; the concept of time as a system of cycles at multiple scales has much broader application (e.g., periodic physiological processes nested within life cycles nested within population cycles) and does not need to be limited to periodic processes with regular period lengths.

Nested periodicity can be represented by hierarchical helices, also referred to as multiply-twisted helices. Conceptually, if a helix strings together repeated cycles of a given period length (e.g. a 24-hour day), then another periodic event (e.g. a 29.5-day lunar orbit) can be integrated by twisting the original helix into a new helix. This is equivalent to wrapping a helix with periods at a lower temporal scale (e.g. days) along one with periods at a higher temporal scale (e.g. months or years). The concept is illustrated in Figure 3 and described with notation in the Methods.

**Figure 3:**
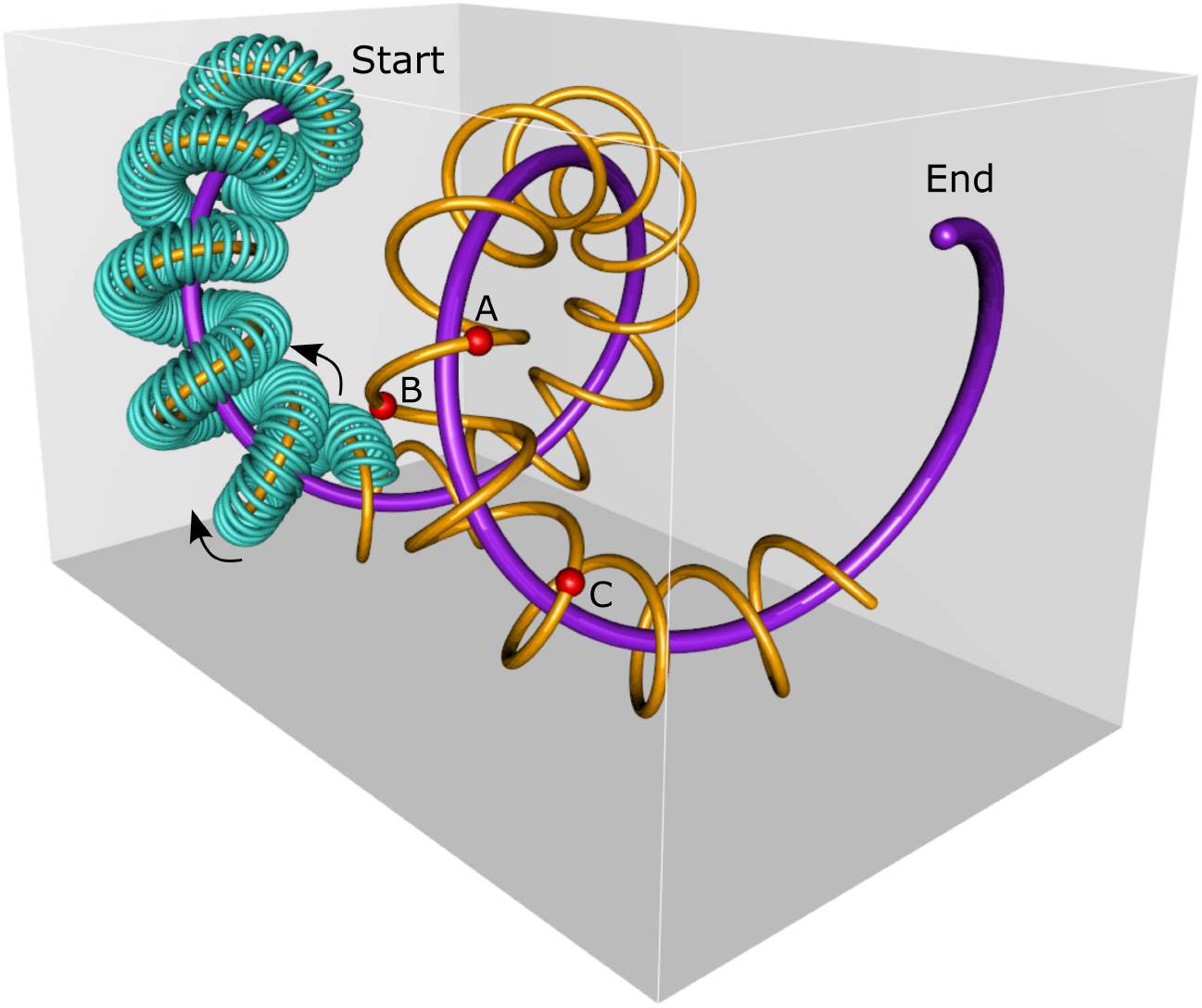
Illustration of the concept of the hierarchical helix, with helix orders indicated by different colors: purple - 1^*st*^ order, yellow - 2^*nd*^ order, and turquoise - 3^*rd*^ order. Three events taking place at different times along the 2^*nd*^ order helix are marked with red spheres. Note that it is the path of the highest-order hierarchical helix (turquoise) that represents temporal progression (direction indicated with arrows), not any of the individual axes delineating the hyperspace. In the case of helices of 2^*nd*^ and higher order, it is possible for a later event (B) to precede an earlier one (A) along any of the major axes.

This process can, in theory, be repeated to reach any desired level of helical hierarchy (see Methods). In practice, the benefits of hierarchical visualization are bound to quickly be outweighed by the increasing complexity and ungainliness of the construct. Here I will only discuss 3^rd^ and lower order helices. It is worth noting that there are physical manifestations of hierarchical helices (e.g., multiply-coiled tungsten filaments, super-helical DNA), and their mathematical and physical properties are being explored^16^,^17^.

### Scale-transcending visualization

Traditionally, patterns in temporal data are represented separately at different time scales, often by collapsing other scales through aggregation or by ignoring that dimension of the data. For example, plots meant to convey seasonal activity levels of an animal may show aggregated daily activity over the course of a year^18^, whereas diel activity plots may project hourly activity data collected over an extended period onto a single (hypothetical) day^18^,^19^, ignoring or collapsing the sequence of multiple days over which observations actually took place. To convey changes in diel activity over the course of a year, data collapsed to a daily scale (e.g., aggregated by hour of the day) can be organized into a series of seasonal snapshots^20^. Multiple scales can be organized in array form, with the dimensions of the array representing the different nested cycles. An example is a paper calendar that arranges days of the week and weeks of the month as rows and column of a matrix, with individual pages for each month representing matrix slices. Similar array structures can be used to represent nested cycles^19^, but they do not capture both periodicity and time continuity. Due to the constraints on hyper-dimensional visualization and the linear vs circular paradox, the aforementioned methods are limited or unable to display extensive temporal data sets at their full detail and reveal patterns potentially interacting at multiple temporal scales.

The hierarchical helix model offers a more general and flexible metaphor for continuous time that allows conceptualization and visualization at multiple temporal scales simultaneously in three dimensions, possibly projected onto a two-dimensional canvas. Both categorical and numerical data that describe conditions or events (and aggregated versions thereof) can be integrated through coloring the observations that make up the helix structure. Alternatively, they can be added as additional visual elements (e.g., points, spheres, or lines) superimposed onto the basic structure.

Periodicity at each scale of the hierarchical helix is made visible through proximity. As such, the approach can reveal not only patterns manifesting at different temporal scales, but also dynamics in such patterns across scales. Temporal data sets that are fine in grain (e.g. hours) and large in extent (e.g. years) are challenging to display in their totality using linear representation. The hierarchical approach thus brings along an added geometric benefit: “coiling” compresses an otherwise unwieldy time series into a much smaller space (achieved through a perspective view of a time spiral by^23^). This is demonstrated in Figure 4, which shows two-year time series of hourly light and temperature data configured as 3^*rd*^ order helices and revealing clear patterns in environmental conditions with both fine temporal grain and large temporal extent. Other environmental information, such as noise and air pollution levels, could be similarly displayed in order to convey their dynamics across relevant cycles for human industry (e.g., diel, weekly, and annual).

**Figure 4:**
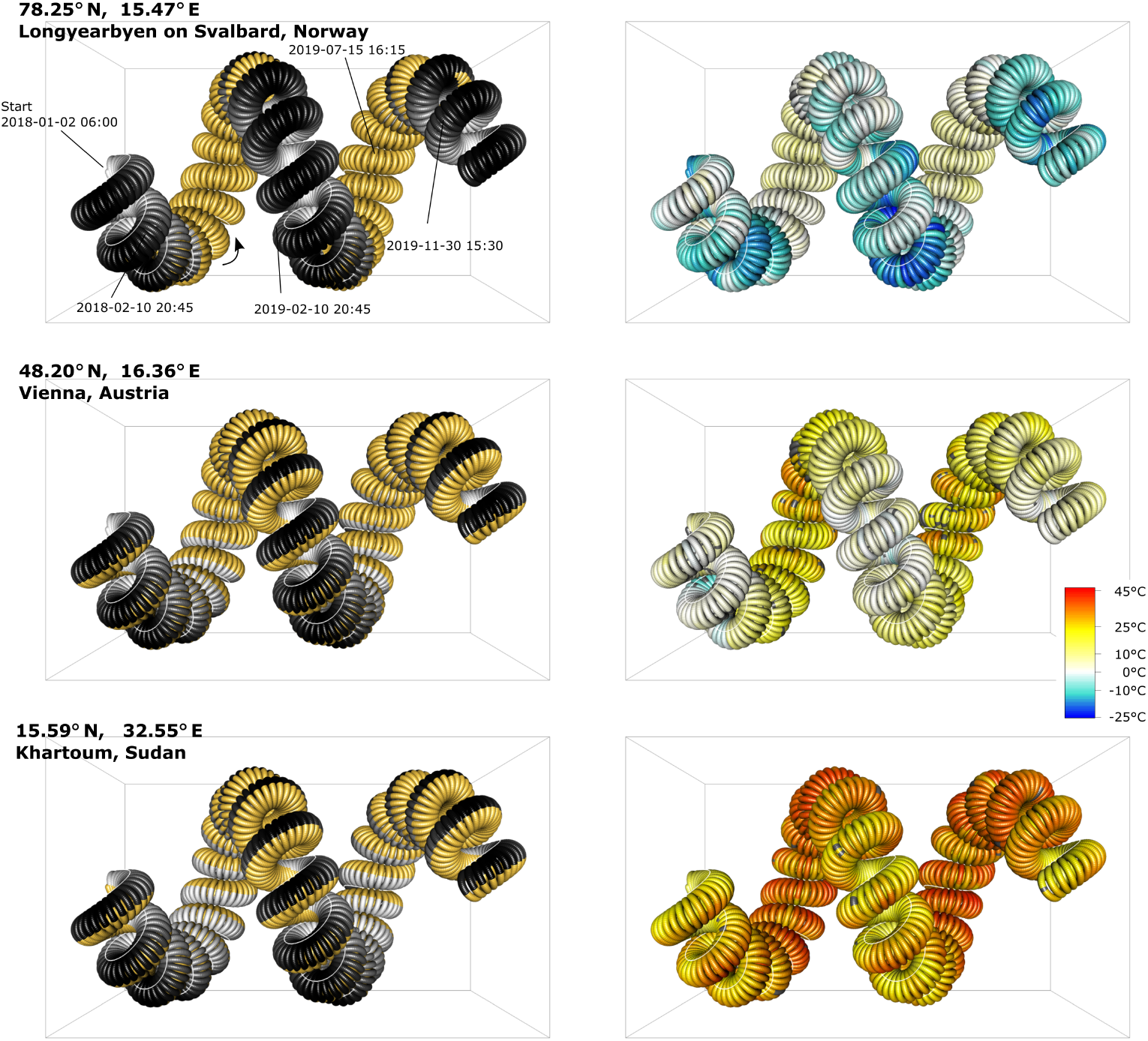
Astronomical light (right) and hourly air temperature (right) at 3 different latitudes (decreasing top to bottom) over two calendar years (2018 and 2019). Colors are rendered onto the same helical construct: year - 1^*st*^ order, lunar month - 2^*nd*^ order, and day - 3^*rd*^ order (see Fig. 3 for orientation). Illumination is shown as daylight (yellow) and night, ranging from full moon (white) to new moon (black). The white line running along the helical structure marks midnight (00:00:00). Sun and moon data were obtained with R package suncalc^21^. Air temperature data were obtained with R package worldmet^22^. Missing values in the temperature plots are shown in gray.

The hierarchical helix model can be extended to allow for non-regular periodic elements. Loop radii and number of return periods at lower temporal scales contained within loops at higher scales can be made functions of intrinsic and extrinsic factors instead of fixed parameters. For example, a speed-up or slow-down in the perceived passage of time, a phenomenon explored in psychology^24^, could be visualized as changing loop diameters or stretching of the helix structure.

One of the disadvantages of projecting a three dimensional object onto a twodimensional display is that, with a fixed perspective, some areas with data are hidden from view^6^. This can be partially solved by displaying helices from different perspectives (Fig. 5) or with rotation (Supplementary Information Figure S1). An immersive visualization approach^25^, is probably the most suitable, as the ability to rotate, shift, and resize the helical construct lets the user find perspectives most conducive for revealing patterns at desired scales. Functions and a vignette with examples for creating such interactive plots are provided at https://github.com/richbi/hHelix. Employing virtual reality applications for exploring hierarchical helices may further mitigate the challenges arising from the complexity of the construct and high-dimensional data^26^.

**Figure 5:**
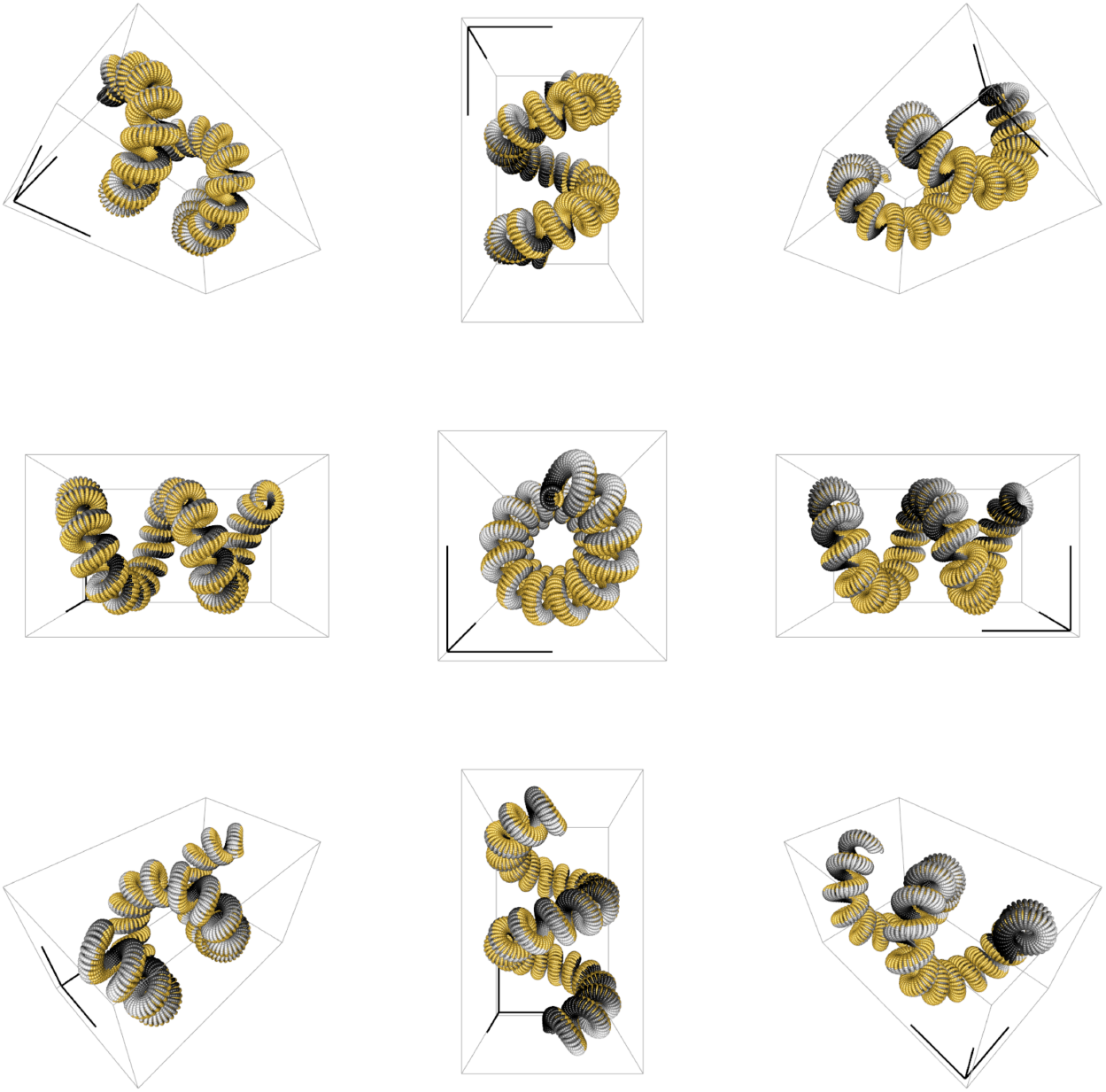
A 3^*rd*^ order hierarchical helix (days - lunar months - years) shown from different perspectives and rendered to display astronomical lighting conditions (daylight [yellow] and night ranging from full moon [white] to new moon [black]). Each peripheral construct can be viewed as a “folded out” version of the central construct; the dark axes lines in each bounding box indicates the same corner, for reference.

Another option for visual exploration is to change the number of orders into which the helix is configured. This can reduce the complexity of the three-dimensional construct, which may in some cases make patterns more pronounced. For example, Figure 6 shows Eurasian badger (*Meles meles*) activity data recorded at automatic wildlife cameras in Norway over a three-year period. Visual pattern in these plots and interactive versions thereof suggest that a) badger activity is lowest during the winter (not surprising, as this species hibernates in cold climates^18^), b) activity is more strictly nocturnal during the winter half-year (when nights are longer), and c) badgers apparently exhibit lower activity during periods with full moon, at least during the winter half-year.

**Figure 6:**
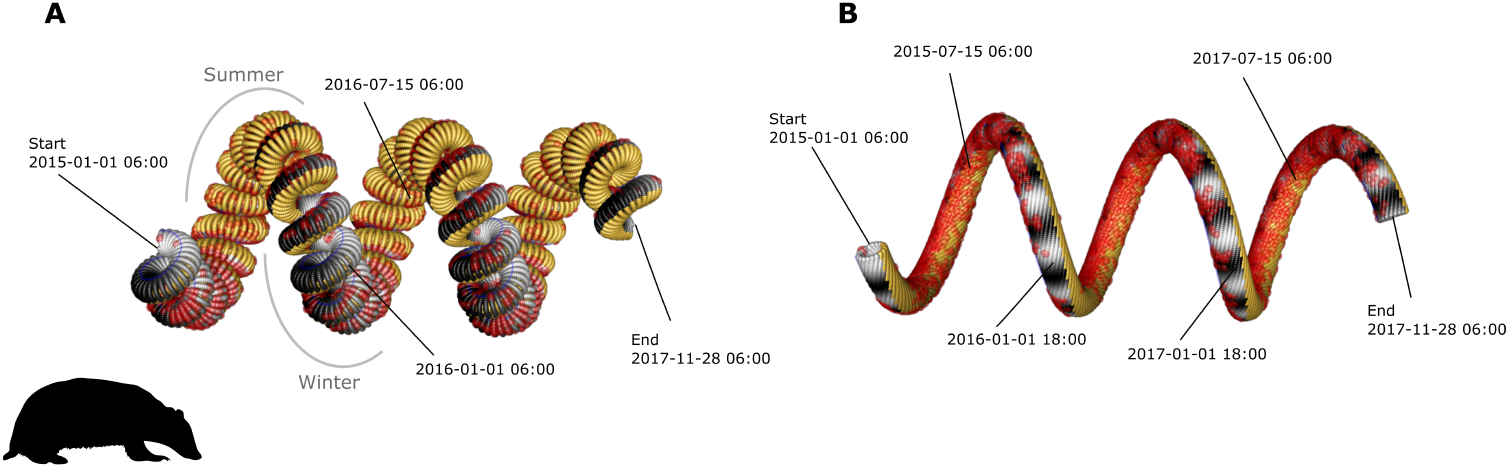
Temporal information projected onto hierarchical helices with orders configured according to different combinations of scales of periodicity: A) 3^*rd*^ order helix with days, months, and years; B) 2^*nd*^ order helix with days and years. Color-rendering of the helical constructs corresponds to patterns of daylight (yellow) and night (ranging from full moon [light] - new moon [dark]). Small red spheres indicate photographic detections (at least one photo during a given hour at a station) of Eurasian badgers (*Meles meles*) at a network of 221 camera traps in southern Norway, deployed during long-term wildlife monitoring by the Norwegian Institute for Nature Research (http://viltkamera.nina.no/). A rotating version of the 3^*rd*^ order hierarchical helix is provided in the Supplementary Materials (Figure S1).

## Concluding remarks

The hierarchical helix is a conceptualization, visualization, and data browsing method - it relies on human intuition. It is not a formal analytical tool for drawing reliable inferences, but can aid in the choice of, and guide, statistical analysis. The self-similar helical model of time can serve as a metaphor and cognitive map for nested periodicity that reconciles the linear-periodic paradox. Technical developments have made interactive and immersive visualization more widely accepted and accessible, which can help overcome the challenges associated with visualizing and browsing complex, hyper-dimensional information. Readers can explore the method with the collection of functions and a vignette available at https://github.com/richbi/hHelix.

## Acknowledgements

I thank J. Odden and project Scandcam at the Norwegian Institute for Nature Research for permission to use the badger camera trap data. I thank J.G.O. Gjevestad and O. Bischof for discussions and C. Milleret and P. Dupont for comments on the manuscript.

## Author contributions

This work was conceived and executed by RB.

## Materials & Correspondence

Correspondence and requests for materials should be addressed to the author.

## Data availability

Data and model code are available at https://github.com/richbi/hHelix.

## Methods

### Hierarchical helix formulation

A single, non-hierarchical helix (1^*st*^ order), can be formulated as a vector 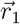 changing with time t in three-dimensions (*x, y, z*).

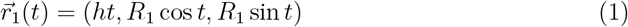

where R_1_is the radius of the loops, *h* the displacement along the x-axes, and the subscript _1_denotes the helix order. Henceforth, it is implied that 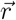 is a function of *t* and “(*t*)” will be omitted for readability.

Following^27^, a 2^*nd*^ order helix 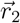 that wraps around 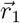 can be constructed by finding the tangent 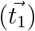, normal 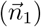, and binormal 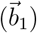 vectors along the trajectory of 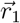.

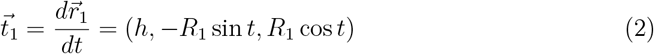

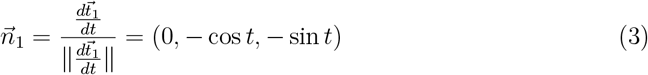

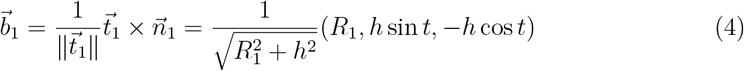

The vector 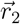 describing the 2^*nd*^ order helix is thus

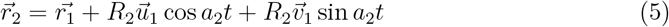

where *R*_2_is the radius of the loops of the 2^*nd*^ order helix and a_2_ is the number of times the 2^*nd*^ order helix wraps around each full loop of the 1^*st*^ order helix.

The aforementioned process is repeated until a helix of desired order *k* is reached.

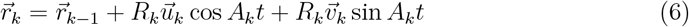

where 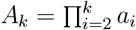 and a_*i*_ is the number of times the helix of order i wraps around each full loop of the helix of order i − 1. A description of doubly and triply-twisted helices is, for example, also provided in^28^.

R^29^ functions for implementing the above vector-based formulation and constructing helices of 3^*rd*^ and lower order are provided at https://github.com/richbi/hHelix, together with utilities for projecting timed data onto interactive plots of hierarchical helices. Use of the functions is exemplified in a vignette, also available at https://github.com/richbi/hHelix.

### Source and preparation of example data

Daily sunrise and sunset schedules, as well as hourly lunar illumination for example locations were obtained with R package suncalc^21^. Hourly air temperature data were obtained using the R package worldmet^22^. Time-stamped records of photographic detections of Eurasian badgers (*Meles meles*) were made available by the Norwegian Institute of Nature Research (https://www.nina.no/). The data were collected as part of a long-term project (Scandcam, http://viltkamera.nina.no/) using motion-triggered cameras mounted to trees in forested areas. The example in this study was limited to a subset of 210 camera sites and three years (2015-2017), consisting of 6 836 photographic detections of badgers. The study area is located in the south of Norway (10.92^*?*^ E, 59.69^*?*^ N) and compasses 18 000 km^2^. For details about the camera trap data collection, see^30^. Photographic detections at each camera trap location were aggregated into hourly values with observations representing at least one detection of a badger at a given station during a given hour. The resulting data from all stations was projected onto the same hierarchical helical structure using the R functions provided at https://github.com/richbi/hHelix to visualize detection intensity over time.

