## Supplementary figures and images for "A self-similar helix as a metaphor for continuous time with nested periodicity"

### Supplementary Figure S1

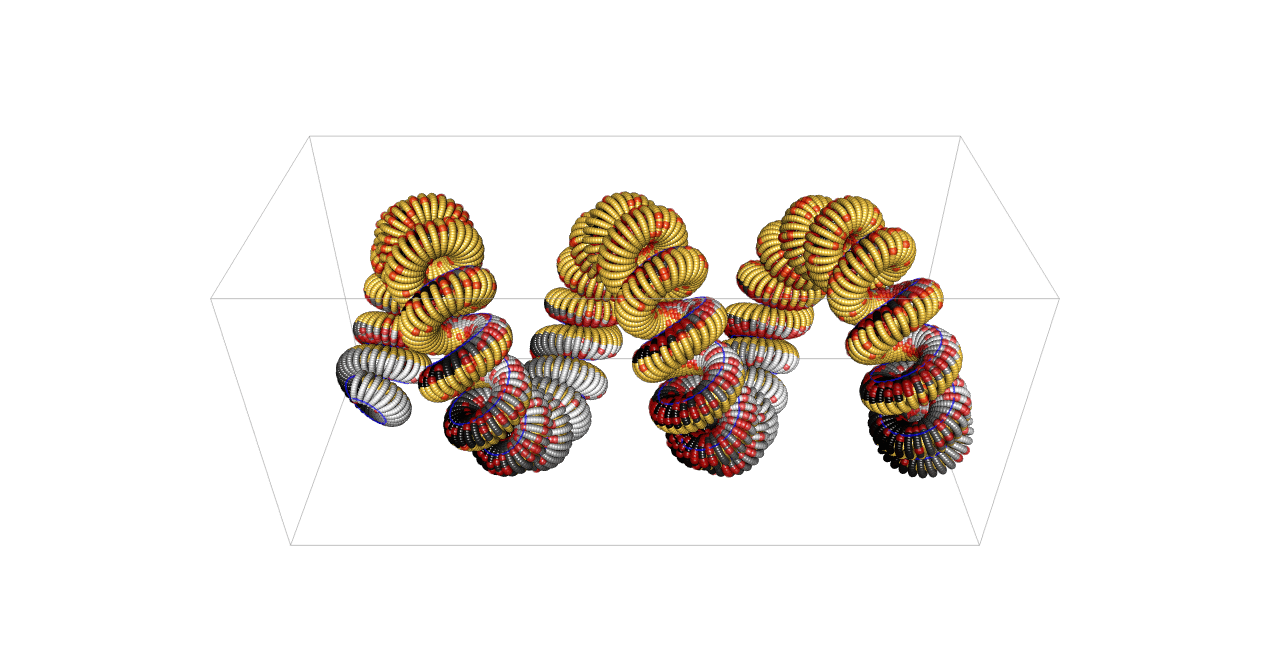
